# A Petri Net–Based Computational Framework for Oxygen-Sensitive Regulation of Pyruvate Metabolism in *Mycobacterium tuberculosis*

**DOI:** 10.1101/2025.09.02.673648

**Authors:** Abhilash, Mohd Asif Siddiqui, Ravindra Kumar Jain, Sangeeta Dayal, Neelesh Kapoor, R. P. Mahapatra

## Abstract

Pyruvate serves as a central hub of cellular metabolism, linking glycolysis with downstream pathways such as the tricarboxylic acid (TCA) cycle, gluconeogenesis, and fermentation. Its metabolic flexibility enables organisms to switch between oxidative and fermentative fates depending on oxygen availability and environmental stress. In this study, a Petri net (PN) model of pyruvate metabolism in *Mycobacterium tuberculosis* H37Rv was developed to capture and simulate these dynamic transitions. Using Snoopy 2.0 for model construction and COPASI for simulation, the framework incorporated key metabolites, cofactors, and enzymatic processes regulating aerobic and anaerobic states. The model demonstrated that oxygen availability acts as a regulatory switch, channeling pyruvate either toward acetyl-CoA for ATP generation under aerobic conditions or toward lactate production under hypoxia to regenerate NAD^+^. Structural validation confirmed boundedness, conservativeness, and deadlock-free behavior, underscoring the robustness of the framework. Sensitivity analyses highlighted enzymatic kinetics as critical determinants of flux distribution and system stability. Collectively, the PN model provides a scalable and biologically relevant computational framework for exploring oxygen-dependent metabolic reprogramming, offering insights into energy adaptation and potential therapeutic targets in pathogenic systems.

## Introduction

Pyruvate is a central metabolite situated at the crossroads of energy metabolism, serving as a critical junction between catabolic and anabolic processes. Its pivotal role in cellular physiology arises from its ability to channel carbon flux into multiple downstream pathways depending on the environmental context and cellular demands (Méndez-Lucio & Medina-Franco, 2017). Under aerobic conditions, pyruvate is predominantly directed toward oxidative decarboxylation, yielding acetyl-CoA that enters the tricarboxylic acid (TCA) cycle to drive ATP production through oxidative phosphorylation (Nelson & Cox, 2021). In contrast, under hypoxic or anaerobic conditions, organisms divert pyruvate toward lactate or ethanol production to regenerate NAD^+^ and sustain glycolysis. Additionally, pyruvate functions as a precursor for biosynthetic processes: it can be transaminated to alanine, carboxylated to oxaloacetate to replenish the TCA cycle (anaplerosis), or serve as a substrate for gluconeogenesis. This biochemical flexibility underscores pyruvate’s centrality in maintaining cellular homeostasis, energy balance, and redox regulation across diverse organisms ranging from prokaryotes to higher eukaryotes.Given its integrative role, perturbations in pyruvate metabolism are closely linked with pathological states such as cancer, neurodegenerative disorders, and infectious diseases, including *Mycobacterium tuberculosis* infections where metabolic adaptability is key for pathogen survival (Eisenreich *et al*., 2010; Heiden *et al*., 2009). Thus, comprehensive models that capture the fate of pyruvate under varying conditions are essential for advancing systems-level understanding and identifying therapeutic opportunities.

In recent years, computational modeling has emerged as a cornerstone of systems biology, offering a means to integrate heterogeneous data and predict emergent behaviors of complex biochemical networks (Kitano, 2002). Unlike reductionist approaches, computational frameworks allow holistic interrogation of metabolic regulation, robustness, and adaptability under dynamic physiological environments. Among the array of modeling paradigms, Petri nets have gained recognition as a powerful formalism for representing biochemical pathways (Heiner *et al*., 2008). Petri nets are bipartite graphs in which metabolites are depicted as *places* and enzymatic reactions as *transitions*, with tokens representing molecular quantities and directed arcs encoding stoichiometric relationships. This dual graphical-mathematical representation enables both qualitative and quantitative analyses, including reachability, invariants, flux distributions, and dynamic simulations.The current study develops a Petri net model of pyruvate metabolism to explore its diverse metabolic fates under aerobic and anaerobic conditions. By mapping the structural and dynamic properties of this network, the model aims to bridge qualitative visualization with system-level analysis. Such an approach provides deeper insights into how organisms reconfigure metabolic fluxes in response to nutrient availability, oxygen tension, and stress signals. Moreover, the framework offers translational potential by identifying critical nodes and pathways amenable to therapeutic intervention, particularly in pathogenic systems where metabolic reprogramming underpins survival and virulence.

## Methodology

To effectively model the key metabolic pathways of *Mycobacterium tuberculosis* H37Rv, pyruvate metabolism was selected as the representative system. This pathway was considered because of its crucial role in bacterial energy production and its importance as a potential target for novel therapeutic interventions (Matsoso *et al*., 2005; Raman *et al*., 2005). The primary aim was to capture sufficient mechanistic detail to allow reliable computational analysis and dynamic simulation of the pathway.For the construction of the models, the Petri Net (PN) formalism was applied. In this approach, metabolites were represented as places, enzymatic reactions were depicted as transitions, and the quantities of metabolites were expressed through tokens (Heiner *et al*., 2008). The entire process of constructing and executing the models was carried out using Snoopy 2.0 (Heiner *et al*., 2012), a software platform designed for hierarchical and hybrid Petri Net modeling. The arcs connecting places and transitions were derived from the stoichiometric matrix to ensure structural consistency and accuracy (Hofestädt& Thelen, 1998). To represent the presence of enzymes without consuming tokens, test arcs were introduced, while source and sink nodes were incorporated to model the inflow and outflow of biochemical species in the cellular environment. This framework provided a robust and scalable basis for simulating pathway dynamics, identifying regulatory bottlenecks, and enabling system-level exploration of the metabolic processes of *M. tuberculosis* (Westergaard *et al*., 2007).

Dynamic simulations were then performed to investigate the time-dependent behavior of metabolites within the modeled pathways under varying physiological conditions. Both aerobic and anaerobic states were explored, with special attention to the differences in ATP availability under these conditions (Eoh & Rhee, 2013). The simulations were carried out using COPASI (Hoops *et al*., 2006), an open-source tool for the modeling and analysis of biochemical networks. The results generated from these simulations included concentration profiles of substrates, intermediates, and end-products, as well as metabolite flux distributions. ATP yield under oxygen-limited conditions was also assessed, and comparisons were made to examine the energy transitions between aerobic respiration and anaerobic glycolysis (Watanabe *et al*., 2011).

Furthermore, sensitivity analyses were conducted to study how changes in enzymatic kinetics, particularly the rate constant (k_2_), affected pathway performance and overall system output (Schilling *et al*., 2000). To ensure biological plausibility and computational reliability, the Petri Net models underwent rigorous validation. They were assessed for boundedness, conservativeness, and deadlock-freedom, ensuring that the system maintained finite metabolite capacity, mass balance, and accessibility of transitions throughout the modeled processes (Heiner *et al*., 2008). Stochastic verification further established the stability and credibility of the computational framework (Blätke *et al*., 2011).

## Results and Discussion

The Petri Net (PN) model developed to simulate pyruvate metabolism provides a mechanistically coherent and phenomenologically robust framework for analyzing metabolic bifurcation under aerobic and anaerobic cellular environments (Figure 1). Configured with 12 places and 3 transitions, the network incorporates central metabolites, cofactors, and enzymes—specifically pyruvate, NAD^+^/NADH, coenzyme A (CoA), lactate, and acetyl-CoA. These elements capture the essential steps of central carbon metabolism, governed primarily by lactate dehydrogenase (LDH; EC 1.1.1.27) in oxygen-limited states and the pyruvate dehydrogenase complex (PDC; EC 1.2.4.1) when oxygen is sufficient (Santos et al., 2019).

**Figure 1:**
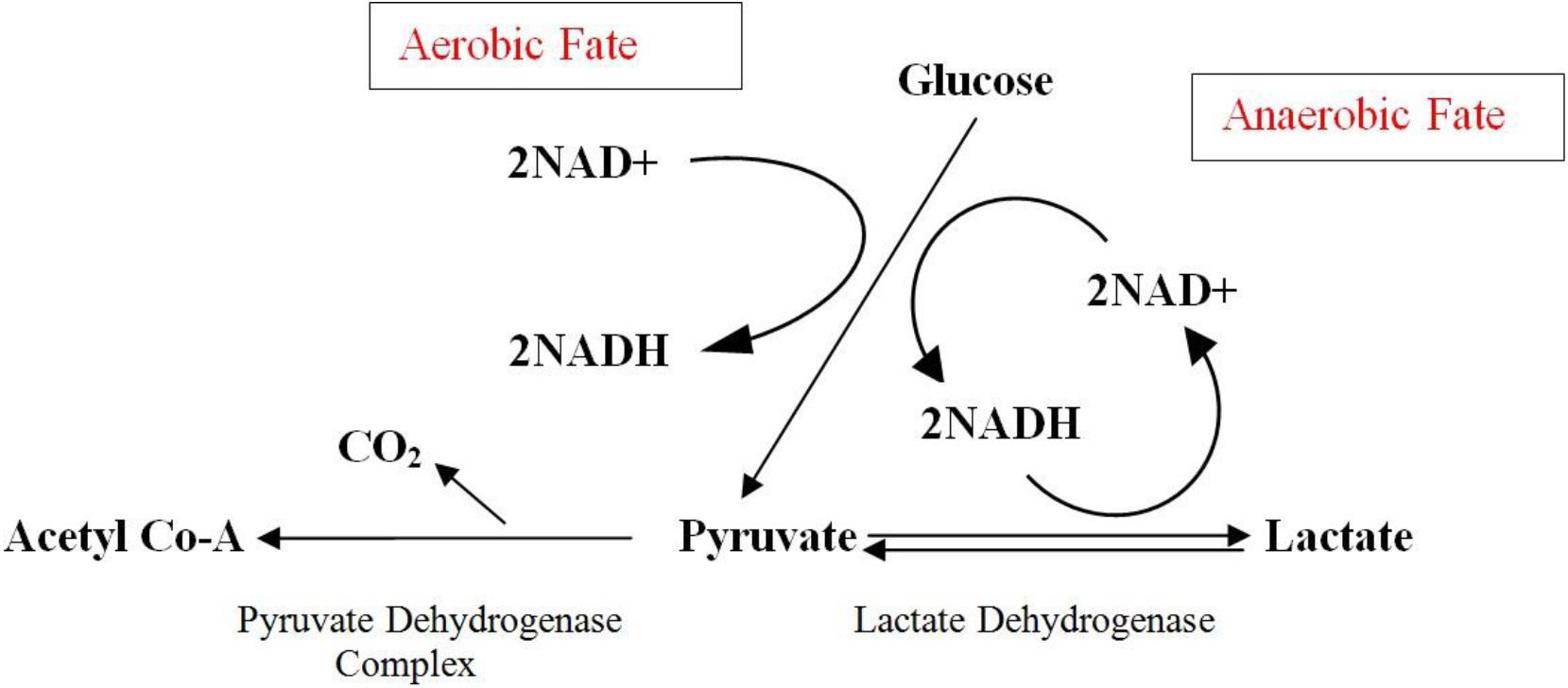
Aerobic and Anaerobic Metabolic Fates of Pyruvate Derived from Glucose.

The PN formalism is particularly suited for metabolic modeling due to its ability to represent metabolites as discrete “places” (biochemical states) and enzymatic processes as “transitions” (reaction steps), thus mapping tokens to molecular concentrations (Heiner et al., 2008). This abstraction maintains close alignment with biochemical processes. PN-based models have been successfully applied to glycolysis (Gilbert et al., 2007), the tricarboxylic acid (TCA) cycle (Blätke et al., 2011), and bacterial energy metabolism (Geistlinger et al., 2022), demonstrating their robustness in capturing complex metabolic dynamics. Within this framework, aerobic metabolism is defined by the oxidative decarboxylation of pyruvate to acetyl-CoA, fueling the TCA cycle and oxidative phosphorylation, whereas anaerobic metabolism channels pyruvate to lactate, regenerating NAD^+^ to sustain glycolytic flux (Eoh & Rhee, 2013; Watanabe et al., 2011). This dichotomy, central to cellular adaptation, has been documented in cancer metabolism (Alberghina et al., 2012) and microbial systems (Matsoso et al., 2005), underscoring the generalizability of PN models across biological contexts.

A key strength of the present PN model lies in its use of *test arcs* to simulate enzyme activity. These arcs enable reactions to proceed without consuming tokens, reflecting the catalytic role of enzymes (Sayah & Chenafa, 2021). Additional structural elements, such as source and sink transitions, ensure metabolite inflow and outflow while maintaining homeostatic balance (Hofestädt & Thelen, 1998). The model begins with pyruvate in a steady-state configuration, and subsequent metabolic trajectories are governed by oxygen availability (Figure 2). Such conditional switching has been shown to strongly influence energy yield and metabolic states in prior computational studies (Schilling et al., 2000; Hoops et al., 2006). When oxygen concentrations exceed ∼20 µM, the PDC is activated, converting pyruvate into acetyl-CoA, NADH, and CO_2_ to support oxidative phosphorylation (Figure 3). Below this threshold, LDH reduces pyruvate to lactate, simultaneously regenerating NAD^+^ to sustain glycolysis (Figure 4). This metabolic reprogramming aligns with physiological scenarios such as muscular exertion, red blood cell metabolism, and ischemic conditions (Schurr & Gozal, 2011; Hochachka & Somero, 2002). The simulations were conducted using Snoopy 2.0 and COPASI, which allowed detailed mapping of temporal and spatial token dynamics (Heiner et al., 2012; Hoops et al., 2006). Structural validation confirmed that the PN system was bounded (finite metabolite pools), conservative (mass balance maintained by P-invariants), and deadlock-free, ensuring biologically realistic behavior (Davidrajuh, 2019; Srinivasan et al., 2017). The model effectively captures the oxidative–fermentative switch in pyruvate metabolism, reflecting foundational principles of mitochondrial respiration (Butow & Racker, 1965). Integration of computational modeling with metabolomics has further demonstrated the regulatory complexity of oxygen-dependent pathways (Buescher et al., 2015; Rizvi et al., 2019). Such multi-omics approaches refine model accuracy and account for context-dependent cellular states (Yizhak et al., 2010). By enabling in silico exploration of oxygen-limited conditions, PN models highlight metabolic plasticity and its role in health and disease (Schuster et al., 2000; Chubukov et al., 2014). This approach also establishes a scalable platform for systems biology, metabolic engineering, and therapeutic applications (Figure 5).

**Figure 2:**
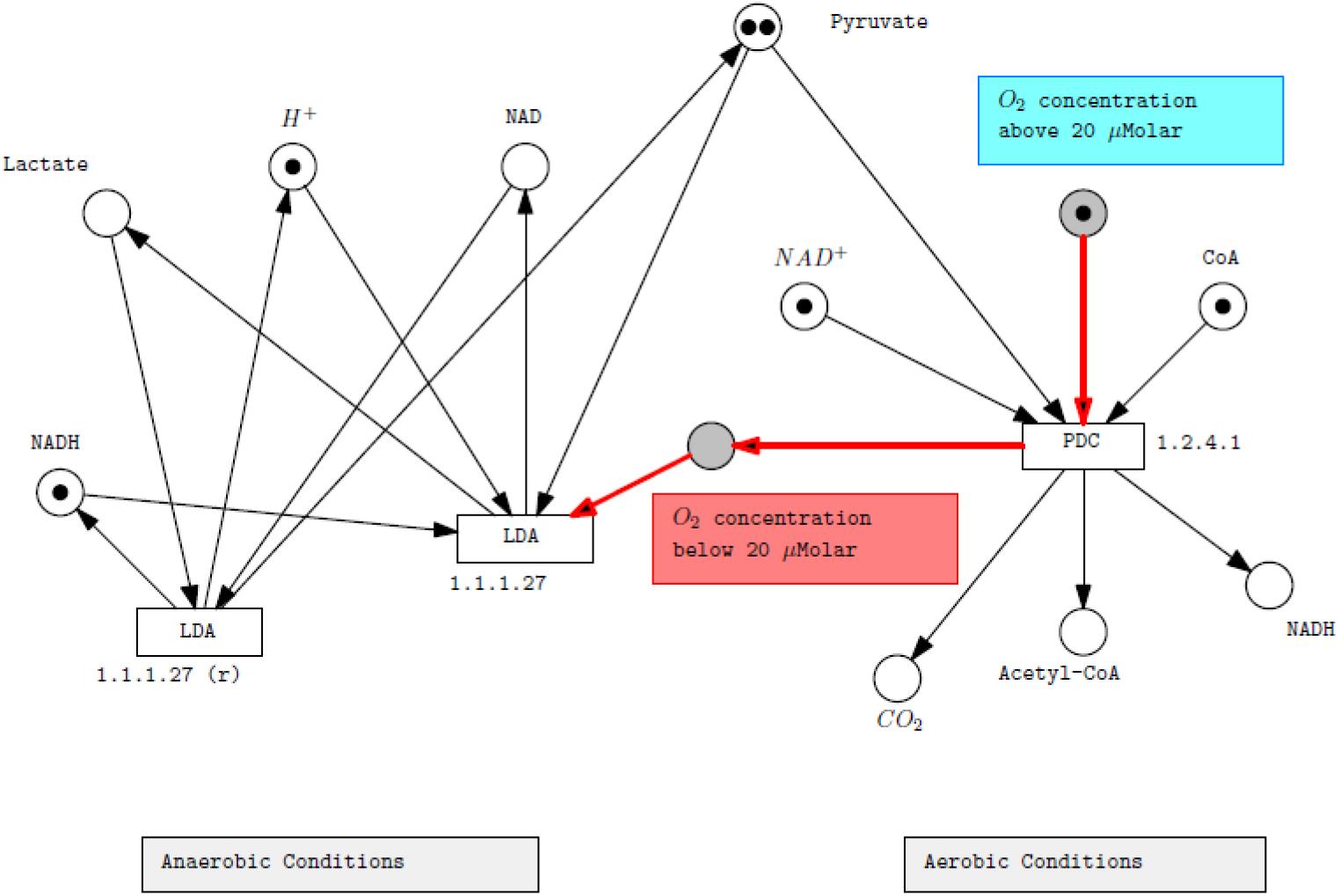
Petrinet model of Oxygen-Dependent Regulation of Pyruvate Metabolism under Aerobic and Anaerobic Conditions (before firing)

**Figure 3:**
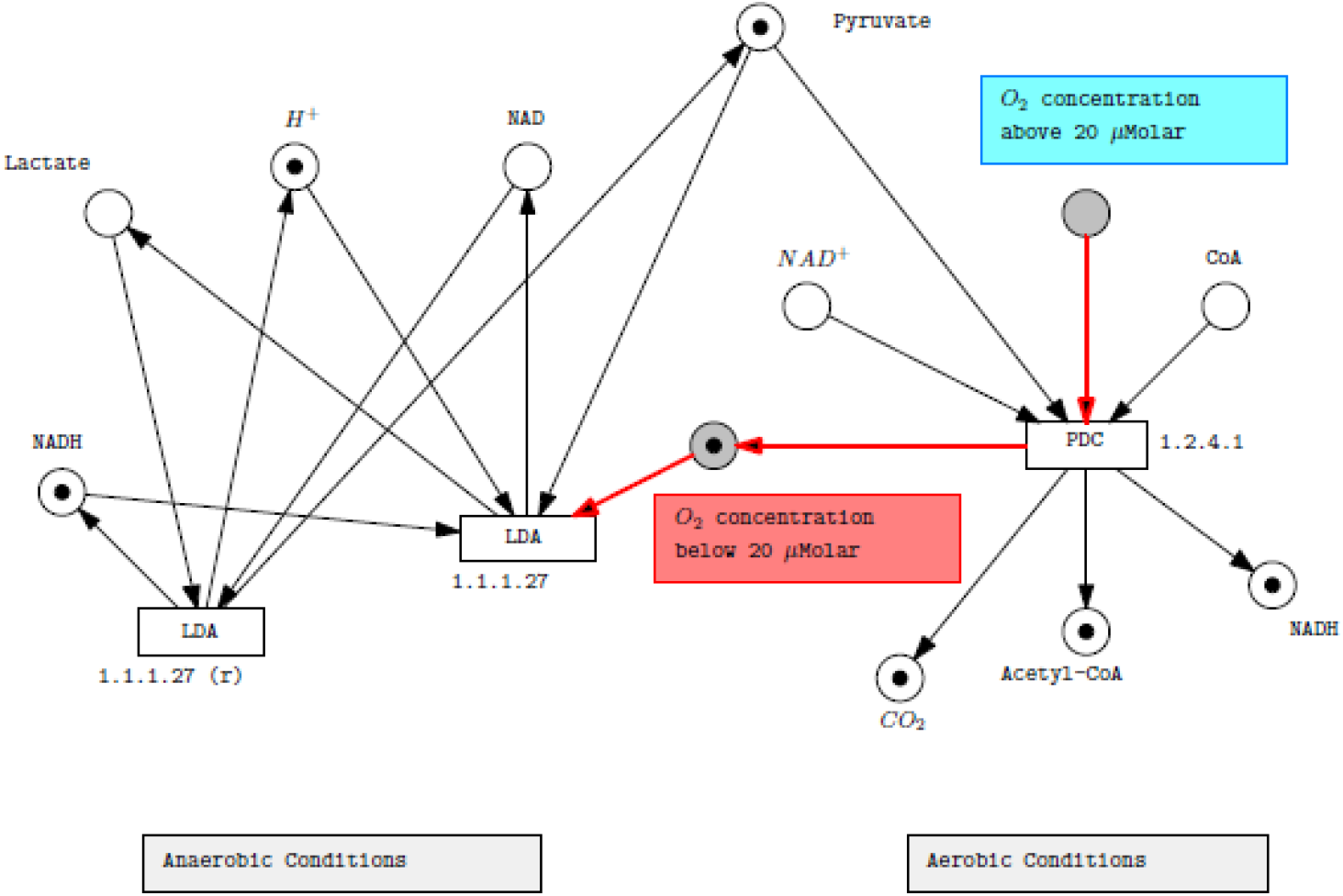
Petrinet model of Enzymatic Regulation of Pyruvate Metabolism under aerobic conditions (after firing)

**Figure 4:**
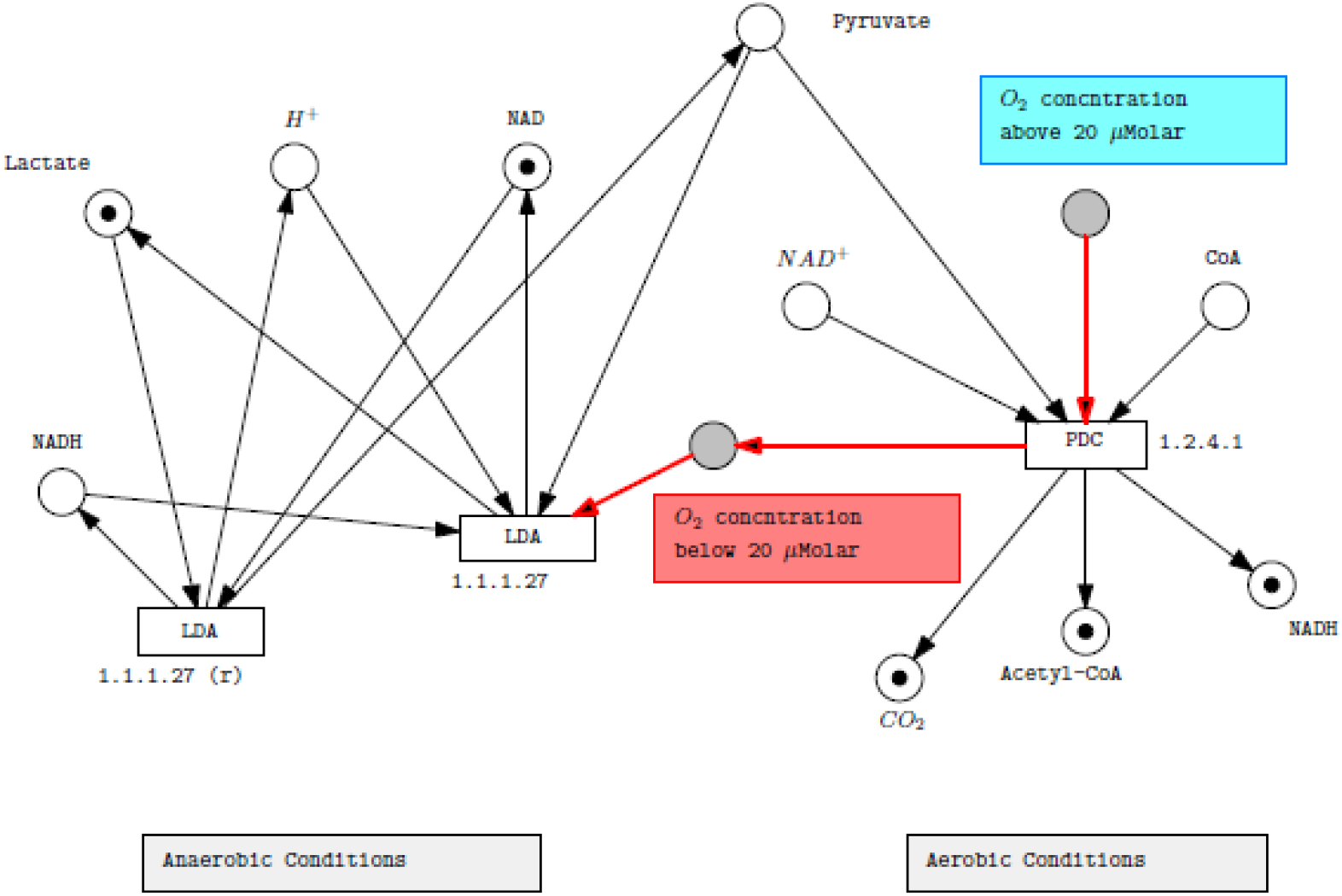
Petrinet model of Oxygen-Sensitive Bifurcationof Pyruvate Metabolism under anaerobic condition (after firing)

**Figure 5:**
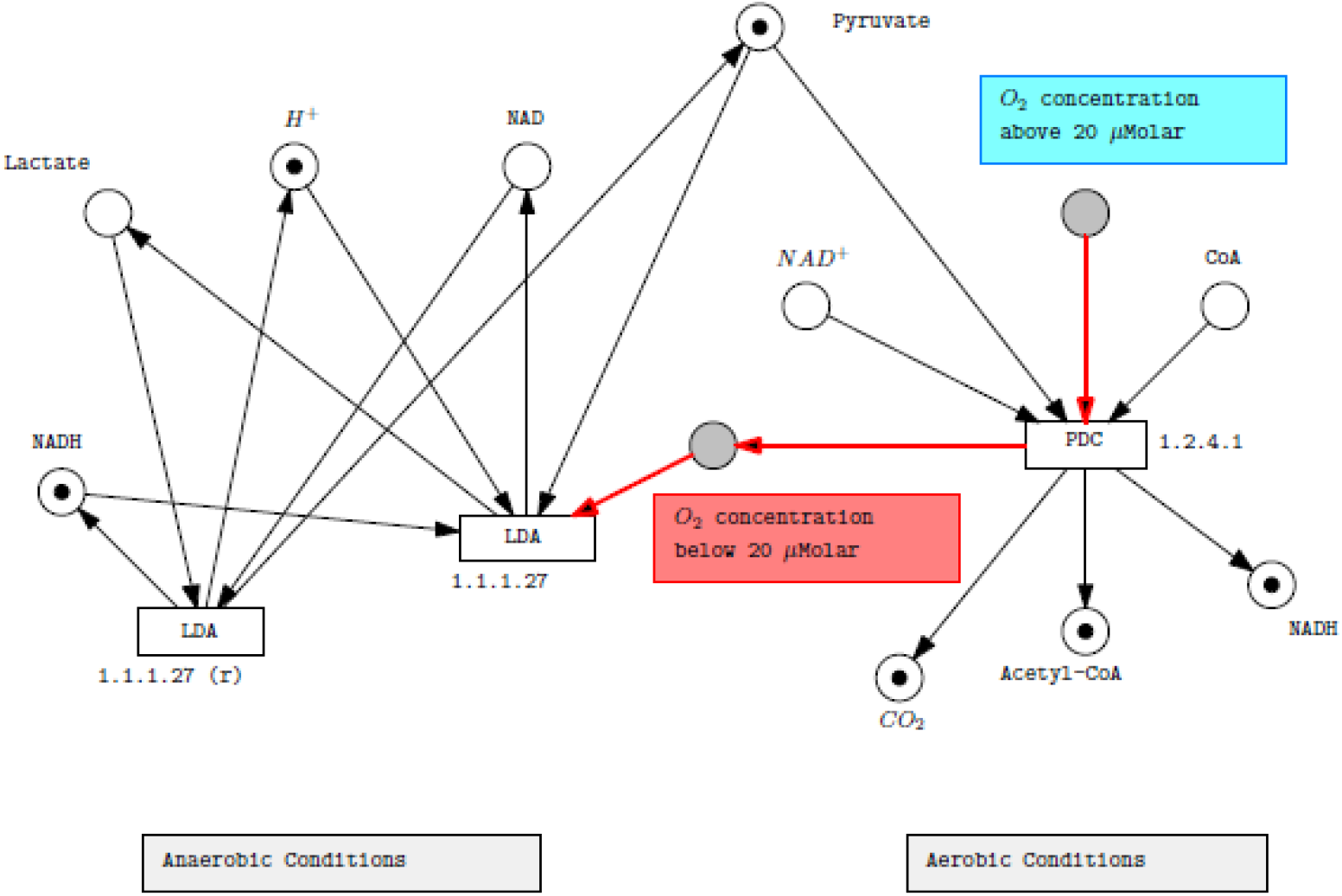
Regulatory Oxygen-Threshold-Driven Switch in Pyruvate Metabolism: A Petri Net Simulation Framework.

Time-resolved simulations underscored the rapidity of metabolic reprogramming: within 60 seconds, cells transitioned from aerobic to anaerobic states, highlighting oxygen as the central determinant of energy metabolism (Ludwig, 2004). Such abrupt oxic–anoxic shifts, observed in cellular and aquatic systems, resemble regime shifts (Bich et al., 2020) and underscore the need to understand dynamic cellular responses to fluctuating oxygen (Bacon & Marsh, 2007). Analysis of metabolite trajectories (Figure 6) revealed a sharp rise in pyruvate concentration within 10 seconds, followed by its depletion as lactate levels increased at ∼15–20 seconds under hypoxia, indicating a switch to glycolysis (Melkonian & Schury, 2019). This mirrors the Warburg effect, where glycolysis persists despite oxygen availability, supporting cell survival under stress (Bose et al., 2020). Simulated oxygen levels declined linearly to zero within 20 seconds (Figure 7), fully enforcing anaerobic metabolism. This triggered compensatory responses, including NAD^+^ regeneration and pathways associated with hypoxia-inducible factor 1 (HIF-1) activation. ATP dynamics (Figure 8) further demonstrated the efficiency gap between oxidative and fermentative metabolism. Aerobic respiration yielded sustained ATP production, whereas anaerobic glycolysis resulted in rapid declines in ATP yield, accompanied by lactate accumulation.

**Figure 6:**
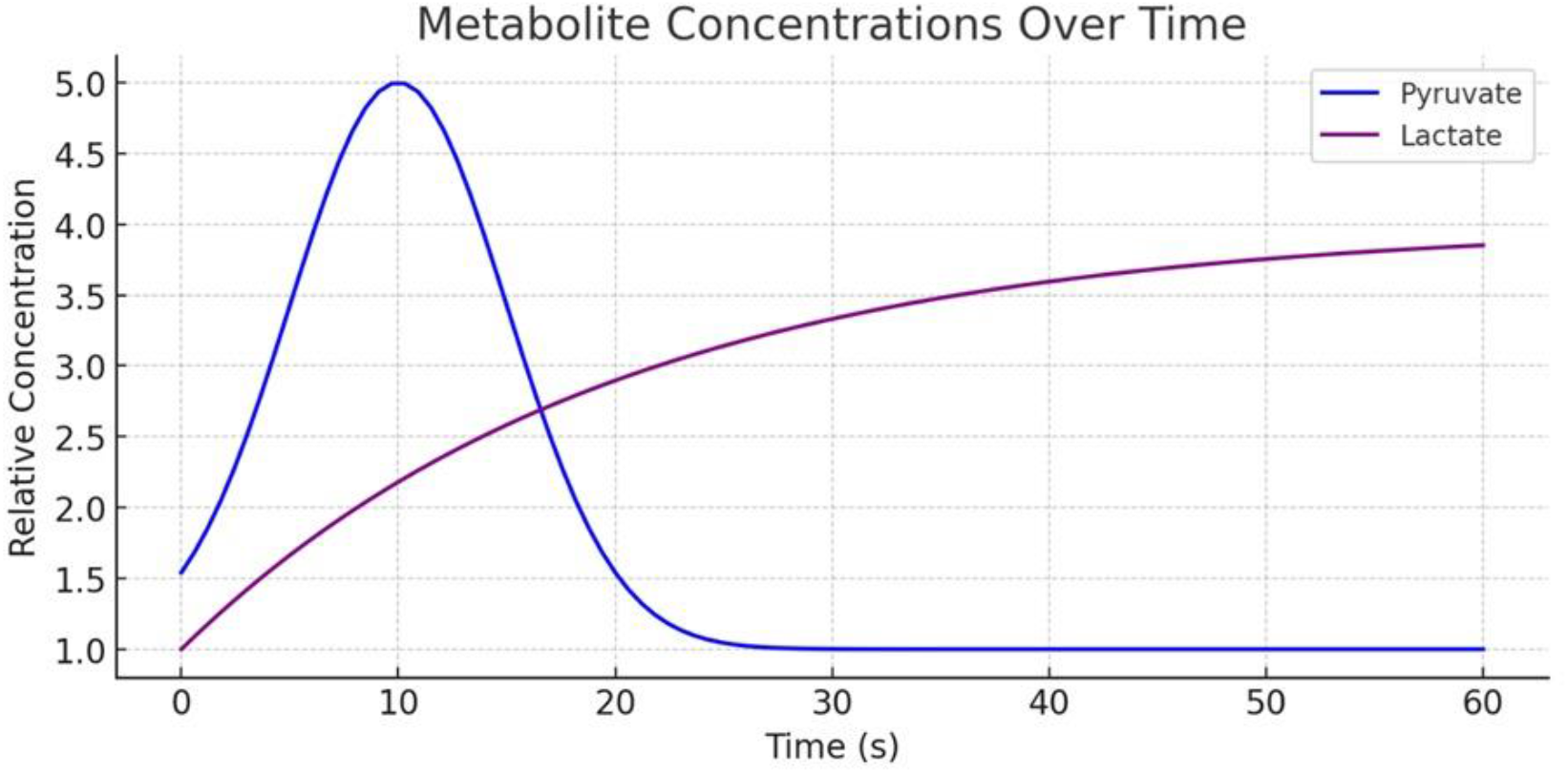
Time-Course Analysis of Glycolytic Metabolites Under Oxygen Depletion.

**Figure 7:**
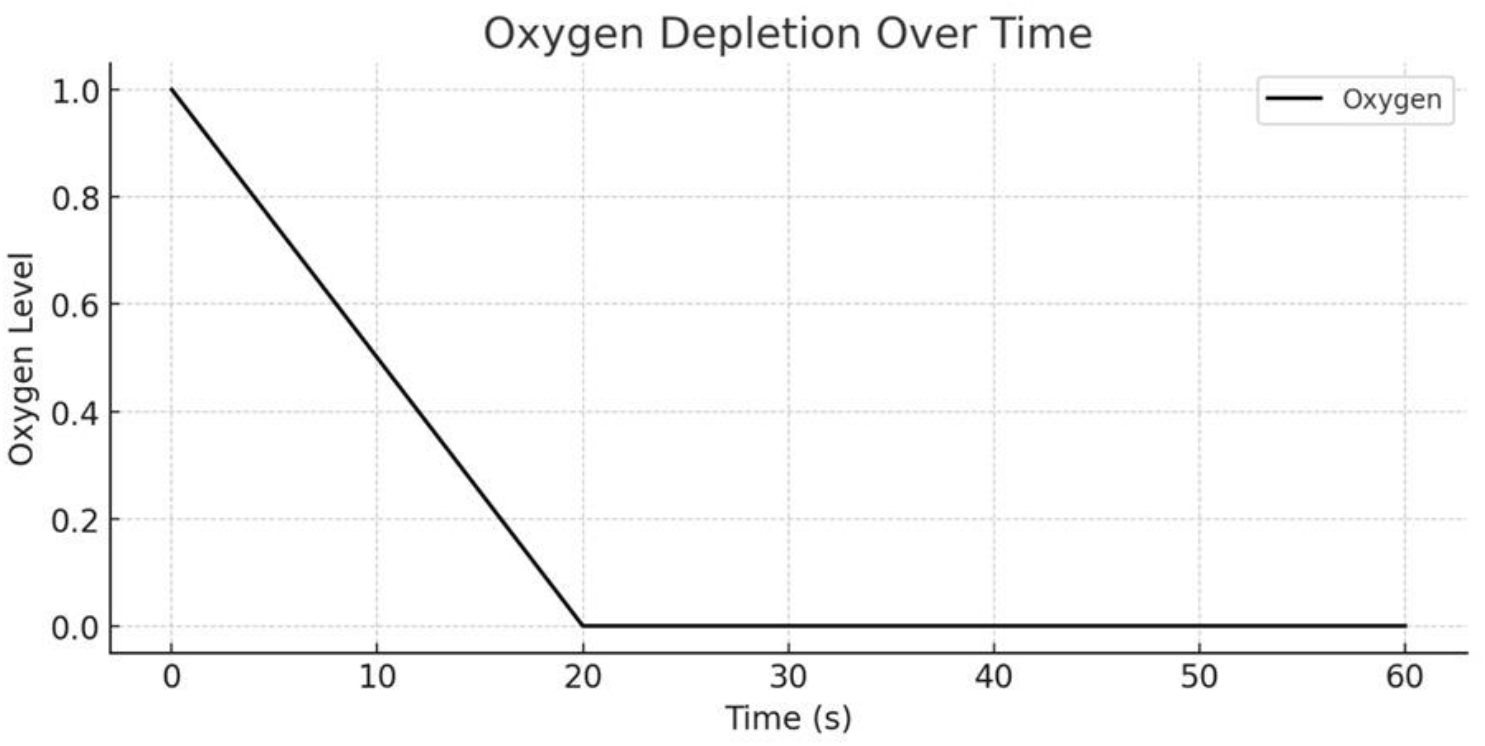
Progressive Oxygen Depletion Simulating Hypoxic Stress Conditions in Cellular Systems.

**Figure 8:**
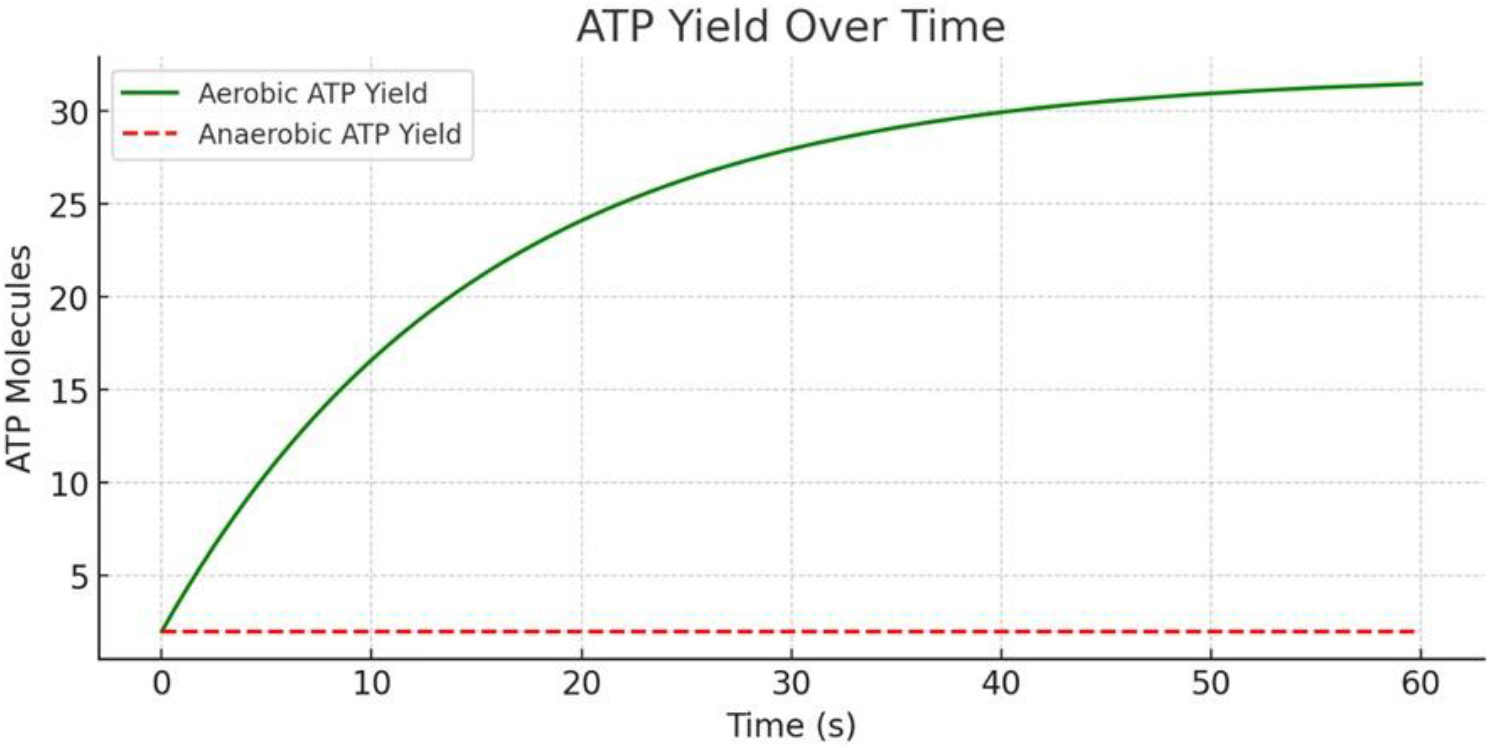
Comparison of Aerobic and Anaerobic ATP Yields During Oxygen Deprivation.

In summary, this PN model provides a rigorous yet flexible framework for simulating oxygen-dependent pyruvate metabolism. The simulations recapitulate fundamental biochemical principles and physiological scenarios, validate structural properties, and highlight the utility of computational models in predicting energy dynamics under hypoxia. Such integrative approaches are valuable for advancing systems-level understanding and for informing therapeutic interventions in ischemia, cancer, and other metabolic disorders.

## Conclusion

The present study establishes a Petri net-based computational framework for understanding the oxygen-sensitive regulation of pyruvate metabolism in *Mycobacterium tuberculosis*. By integrating structural analysis with dynamic simulations, the model successfully captured the bifurcation between aerobic and anaerobic metabolic fates, validated through boundedness, conservativeness, and deadlock-free properties. The findings reinforce pyruvate’s role as a pivotal metabolic switch and highlight the significance of oxygen thresholds in dictating cellular energy strategies. Importantly, the PN approach offers not only a mechanistically interpretable representation of metabolic pathways but also a versatile tool for systems-level exploration. This framework has potential translational applications in metabolic engineering, drug discovery, and therapeutic modeling, where targeting oxygen-dependent metabolic adaptations may provide novel strategies against tuberculosis and related infectious diseases.

